## Supplemental Materials for "General mechanisms of task engagement in the primate frontal cortex"

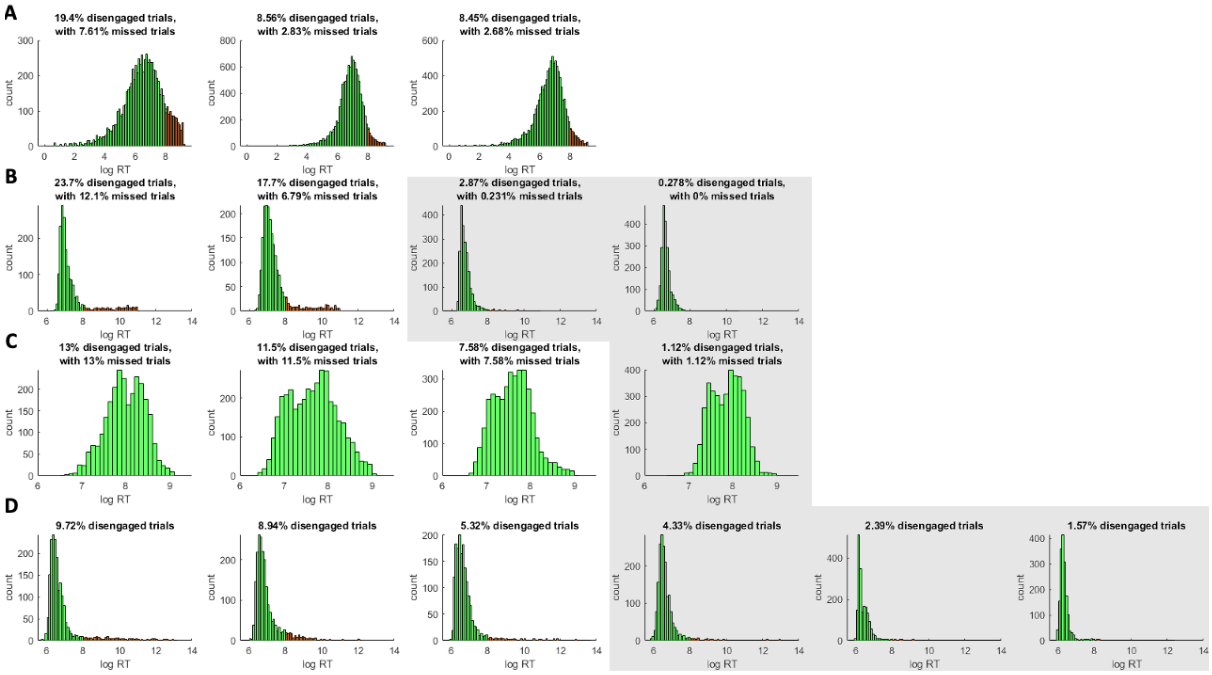


*Figure S1.* ***Response time histograms.*** For each monkey, we binarized trials into engaged (green) and disengaged trials (red). Trials were coded as disengaged if the monkey responded slower than 3s (log(3000ms) ≈ 8) or did not respond at all (missed trials). Animals that had disengaged on less than 5% of trials were excluded from the study (grey background) (**A**) Shows the data from Jahn et al.. In this experiment a trial timed out if the animal did not respond within 10s. The percentage of 'disengaged trials' in the panel title also contain these 'missed trials'. For monkey 1 we had data from 42 sessions, with an average session duration of 83.22 min (std 13.84 min), for monkey 2 40 sessions (average length 64.04 min; std 15.67 min), and for monkey 3 38 sessions (average length 94.22 min; std 12.16 min) (**B**) shows the data from Bongioanni et al.. Here a trial times out if the animal did not respond within 60s. 2 of the 4 animals were excluded from the study as they did not have enough disengagement trials to qualify. For the 2 included animals, we had 12 sessions for monkey 1 (average length 93.08 min; std 14.99 min) and 12 sessions for monkey 2 (average length 81.77 min; std 14.10). (**C**) shows the data from Khalighinejad et al.. In this task, unlike the other 3 tasks, animals had an incentive to delay their response and respond later in a trial to potentially obtain more reward. As such, we did not count trials with RT > 3s as disengaged trials for this task, and only used the missed trials in which the animals did not respond at all to code disengaged trials. 1 animal had to be excluded because it did not have enough overall disengaged trials. For the 3 included animals, we had 23 sessions for monkey 1 (average length 46.13 min; std 4.63 min), 23 sessions for monkey 2 (average length 48.10 min; std (3.76 min), and 21 sessions for monkey 3 (average length 43.46; std 3.37 min). (**D**) shows the data from Grohn et al.. In this task trials never timed-out if the animal did not respond, but instead the stimuli stayed on the screen until the animal re-engaged with the task. As such, there are no missed trials for this task. 3 of the 6 animals had to be excluded because the total number of disengaged trials was less than 5%. For the 3 included animals, we had 13 sessions for monkey 1 (average length 18.90 min, std 6.10 min), 11 sessions for monkey 2 (average length 23.23 min; std 7.38 min) and 12 sessions for monkey 3 (average length 38.97 min; std 14.62 min).

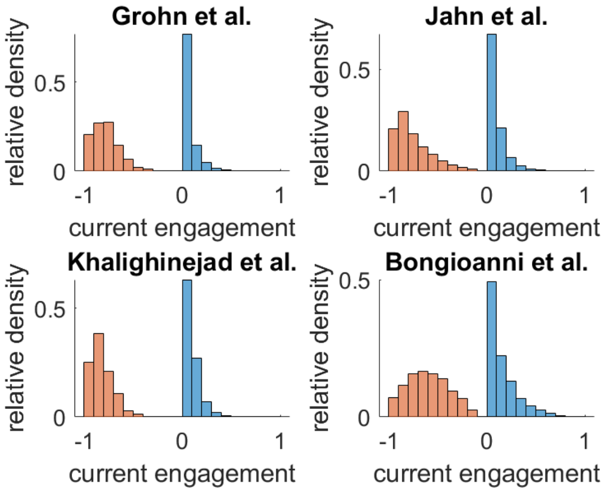


*Figure S2.* Density of current engagement (CE) split by whether the animals were engaged (blue) or disengaged (orange) on a trial. The density of CE for engaged and disengaged trials are normalized separately. For all trials, a value close to 0 indicates that a trial was well predicted by the task-specific regression model. Because CE is the residual of a logistic regression model that predicts engagement with the task, a value closer to 1 indicates that the model predicted a disengagement, but the animal did not disengage. By contrast, a value close to -1 indicates that that the model predicted the animal would be engaged but the animal disengaged on this trial.

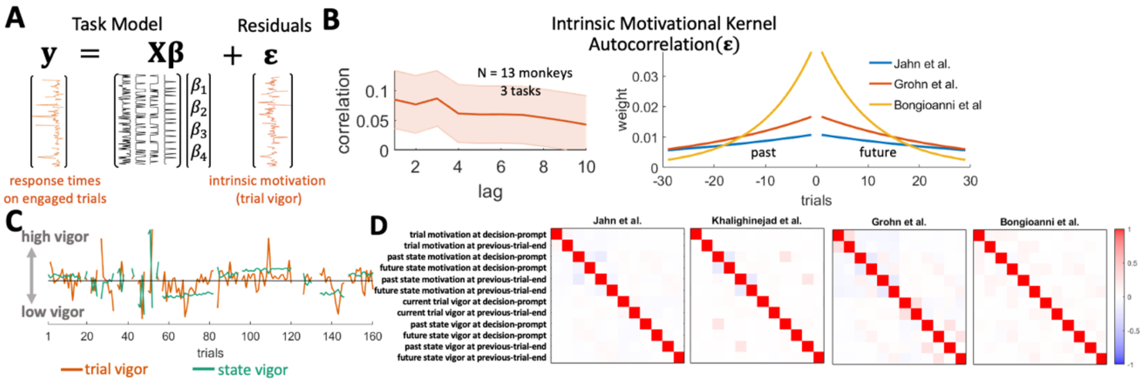


*Figure S3.* ***Vigor as indexed by RTs and whole-brain GLMs.*** (**A**) Just as for task disengagement and engagement as shown in Figure 1, we also constructed separate regression models for the RTs on each task. By regressing out the task here, we were again left with an estimate of trial vigor. (**B**) The residuals of our regression models for RTs are autocorrelated over trials (left; significant for lags < 8). We also again fitted exponential kernels to the residuals of each task (right). (**C**) Smoothing the residuals using these kernels gave us an estimate of state vigor. This estimate of state vigor is distinct from the estimates of task engagement illustrated in Figure 1 as it is based on the residual of the RT regressions and not on the engagement/disengagement regressions. (**D**) Correlations of all regressors in the convolutional models used in the whole brain analysis. Our estimates of the current engagement (CE) and general engagement (GE) (Fig 1) and vigor (this Fig) are not correlated and thus capture different aspects of motivation. We also show the state regressors split up by the past and future state, which can be obtained by using only half of the kernels (i.e. the half directed towards the past or future). By splitting up the state regressors this way we can combine them again on the contrast level to obtain the overall motivational states. We can, however, also subtract them on the contrast level to examine potential differences between past and future levels of vigor.

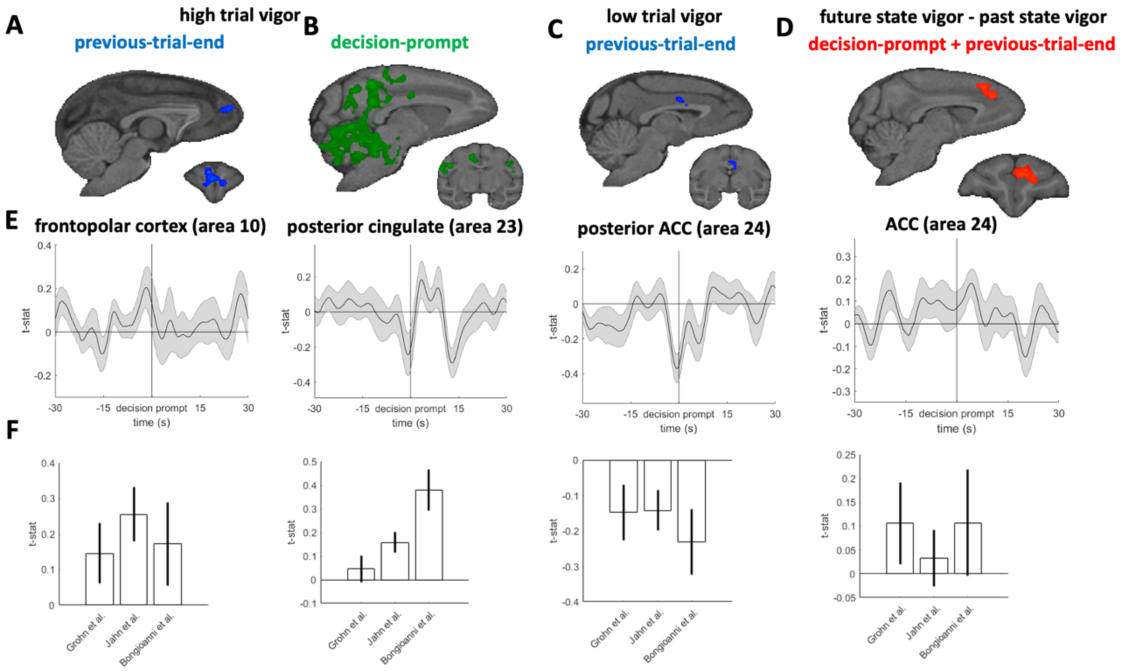


*Figure S4*. ***Transient effects of vigor.*** (**A**). At the end of the previous trial we find activity in the frotopolar cortex when the current trial has high trial vigor. (**B**) However, at decision-prompt of the trial itself high trial vigor is associated with activity in motoric areas in posterior cingulate and cerebellum. (**C**) Low vigor trials are preceded by activity in posterior ACC. (**D**) We also observed activity in ACC/preSMA when animals have higher state vigor in the future than in the past. **(**E) To examine how extended these effects are we visualize their timecourses in ROIs we placed. Unlike the more sustained effects of task engagement as indexed by engagements/disengagements (Fig 2 and 3), effects of vigor are more contained to the current trial. (**F**) Splitting the effects within the ROIs up by task confirms that the signs of the effects are consistent across different tasks.

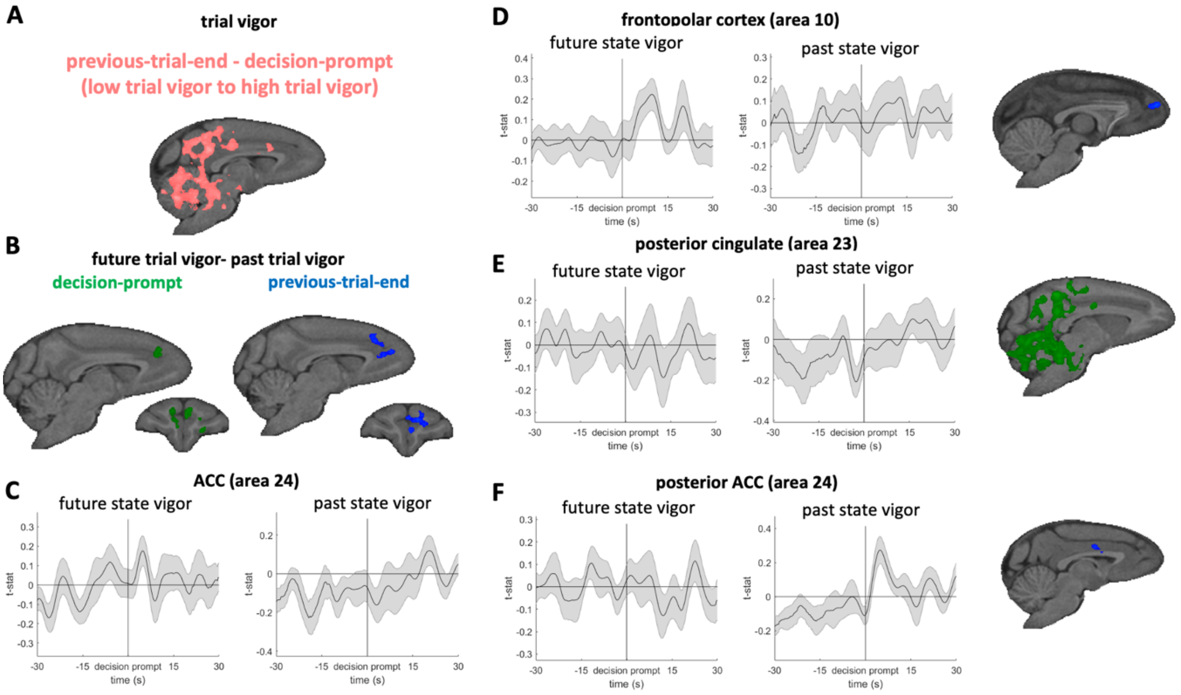


*Figure S5*. ***Transient vigor effects and future and past vigor timecourses.*** (**A**) To examine whether effects of trial vigor were transiently different between previous-trial-end and decision-prompt we examined a contrast of the difference in activity between the two time-points in our tasks. The whole-brain results show the same regions that were found when examining the previous trial end and decision-prompt separately (Fig S4BC). (**B**) When splitting up the activity we found relating to a higher future state vigor than past state vigor (Fig S4D) by decision-prompt and previous-trial-end, we observe the same regions as in Fig S4D. (**C**) Splitting up the timecourse extracted by future and past state vigor shows that activity related to future state vigor is above baseline, while activity for past state vigor is below baseline. (**D-F**) We also examined the activity in all other vigor-related ROIs for effects of past/future state vigor. (**D**) In frontopolar cortex we found increased activity after decision-prompt if animals had more vigor in the future. (**E**) In posterior cingulate we found activity preceding decision-prompt if the animals previously were in a low-vigor state. (**F**) In posterior ACC we found increased activity after decision-prompt if the animals previously were in a high-vigor state.

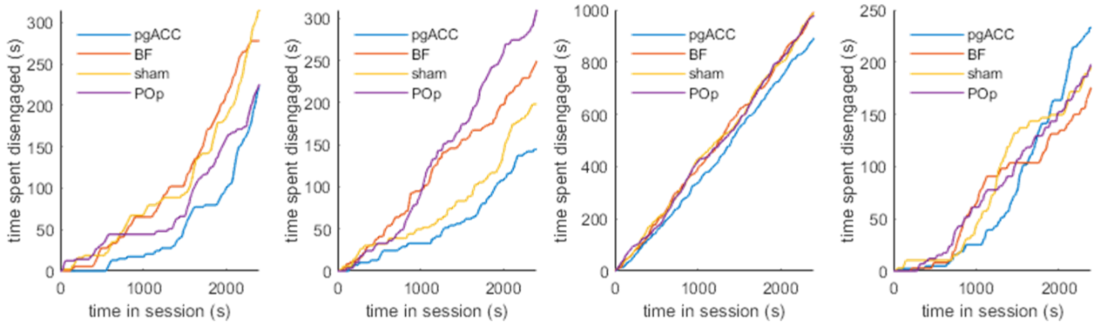


*Figure S6*. ***Time spent engaged after TUS split up by animal.*** The same analysis as in Fig 4C but split up for each of the four animals. The effect of more engagement early on can be seen in each animal.

| Cluster Index | Z | F99 coordinates x | y | z | # voxels | peak description | cluster description |
|---|---|---|---|---|---|---|---|
| 4 | 5.14 | 27.2 | -1.51 | -10.6 | 19261 | right ant STS | right temporal pole, right insula, bilateral striatum, bilateral area 11, 12 and 13, and area 25, 32, and 14 |
| | 4.56 | 27.7 | -3.02 | -9.05 | | right ant STS | |
| | 4.33 | 16.1 | 17.1 | 8.05 | | post central OFC | |
| | 4.22 | -0.503 | 6.54 | -0.503 | | ventral striatum | |
| | 4.2 | 6.54 | 18.6 | 7.55 | | ant central OFC | |
| | 4.17 | 22.6 | 2.52 | -15.1 | | right ant STS | |
| 3 | 4.28 | 29.2 | -16.6 | 5.03 | 3923 | l. mid STS | superior temporal sulcus |
| | 4.2 | 23.6 | -16.1 | 0 | | l. medial mid STS | |
| | 4.11 | 25.2 | -18.1 | 0.503 | | l. medial mid STS | |
| | 3.99 | 28.2 | -7.04 | 7.04 | | l. lateral sulcus | |
| | 3.95 | 24.1 | -12.6 | -5.53 | | l. mid STS | |
| | 3.81 | 27.2 | -3.02 | -1.01 | | l. lateral sulcus | |
| 2 | 4.32 | -21.1 | -4.02 | -1.01 | 2251 | r. lateral sulcus | left insula extending into left claustrum |
| | 4.15 | -18.6 | -2.01 | -1.51 | | r lateral sulcus | |
| | 4.06 | -21.6 | -12.1 | -8.05 | | r medial midSTS | |
| | 4.06 | -22.6 | -4.53 | -4.53 | | r lateral sulcus | |
| | 3.95 | -22.6 | -12.6 | -8.05 | | r mid STS | |
| | 3.95 | -18.6 | -9.05 | 1.01 | | r lateral sulcus | |
| 1 | 5.19 | 18.1 | -8.55 | 19.1 | 1933 | r central sulcus | right central sulcus |
| | 4.67 | 11.6 | -12.6 | 20.1 | | r central sulcus | |
| | 4.45 | 13.6 | -10.1 | 17.6 | | r central sulcus | |
| | 4.42 | 17.1 | -9.56 | 17.6 | | r central sulcus | |

| | | | | |
|---|---|---|---|---|
| 4.36 | 11.6 | -14.1 | 21.1 | r central sulcus |
| 4.34 | 15.6 | -9.56 | 19.1 | r central sulcus |

*Table S1*. ***Significant neural clusters for CE at previous-trial-end and decision-prompt combined.***

| Cluster Index | Z | F99 coordinates x | y | z | # voxels | peak description | cluster description |
|---|---|---|---|---|---|---|---|
| 4 | 4.65 | 4.02 | 10.1 | 4.02 | 15951 | r ventral striatum | right Striatum, right OFC (area 13), frontopolar cortex (10), ACC (32 and 24), SMA and preSMA |
| | 4.54 | 6.04 | 14.1 | 21.1 | | r arcuate sulcus | |
| | 4.51 | 7.55 | 26.7 | 8.55 | | r ant. OFC | |
| | 4.47 | -1.01 | 21.1 | 13.6 | | ant cing. sulcus | |
| | 4.47 | 1.01 | 5.03 | 9.56 | | mid cing. gyrus | |
| | 4.47 | 11.6 | 18.1 | 14.1 | | r principle sulcus | |
| 3 | 6.52 | 11.1 | -13.1 | -11.6 | 10981 | r hippocampus | medial temporal lope, hippocampus and entorhinal cortex |
| | 6 | 11.1 | -11.6 | -12.1 | | r hippocampus | |
| | 5.06 | 24.1 | -6.04 | 0.503 | | r lateral sulcus | |
| | 4.99 | 22.1 | -10.1 | 2.52 | | r lateral sulcus | |
| | 4.99 | 24.1 | -7.54 | -2.51 | | r lateral sulcus | |
| | 4.93 | 26.2 | -21.1 | 13.6 | | r temporo parietal area | |
| 2 | 4.86 | -22.1 | -5.53 | -4.02 | 3822 | l lateral sulcus | left superior temporal lobe extending into left insula |
| | 4.82 | -22.1 | -7.04 | -5.03 | | l lateral sulcus | |
| | 4.79 | -11.6 | 4.02 | -6.54 | | l ventral temp pole | |
| | 4.63 | -20.6 | -4.02 | 2.01 | | l lateral sulcus | |
| | 4.63 | -14.1 | 5.03 | -6.04 | | l ventral temp pole | |
| | 4.61 | -22.6 | -7.54 | -2.51 | | l lateral sulcus | |
| 1 | 5.02 | 16.6 | 7.04 | -9.05 | 3177 | r ventral temp pole | right superior temporal lope extending into right insula |
| | 4.99 | 18.1 | 6.04 | -7.54 | | r ventral temp pole | |
| | 4.89 | 17.1 | 7.04 | -8.05 | | r ventral temp pole | |

| | | | | |
|---|---|---|---|---|
| 4.83 | 12.1 | 8.55 | -2.01 | r posterior OFC |
| 4.73 | 23.6 | 4.53 | -8.55 | r rostral STG |
| 4.41 | 17.6 | 8.05 | -11.6 | perirhinal cortex |

*Table S2*. ***Significant neural clusters for GE at previous-trial-end and decision-prompt combined.***

| Cluster Index | Z | F99 coordinates x | y | z | # voxels | peak description | cluster description |
|---|---|---|---|---|---|---|---|
| 3 | 6.56 | 9.05 | -11.6 | -10.6 | 32606 | r parahippo | right striatum, area 25, anterior area 24, area 32, right area 13, right hippocampus, extending into right temporal pole, right amygdala, right insula |
| | 6.26 | 22.6 | -9.05 | 1.51 | | r lateral sulcus | |
| | 6.06 | 24.6 | -14.6 | 2.52 | | r STS | |
| | 5.72 | 25.7 | -20.6 | 13.6 | | r lat parietal gyrus | |
| | 5.63 | 13.6 | 8.55 | -2.01 | | r posterior OFC | |
| | 5.58 | 10.6 | -10.1 | -11.1 | | r medial temp cortex | |
| 2 | 5.23 | -13.1 | 10.1 | -0.503 | 4643 | l posterior OFC | left insula, left temporal pole |
| | 5.02 | -10.6 | 3.52 | -5.53 | | l ventral temp pole | |
| | 4.95 | -14.1 | 7.04 | -6.04 | | l ventral temp pole | |
| | 4.84 | -15.1 | 5.53 | -4.53 | | l insular cortex | |
| | 4.66 | -13.6 | 8.05 | -3.02 | | l posterior OFC | |
| | 4.63 | -7.04 | 1.01 | -8.55 | | l ventral temp pole | |
| 1 | 6.78 | -21.6 | -2.01 | -2.51 | 3696 | l lateral sulcus | left lateral sulcus |
| | 5.84 | -22.1 | -6.54 | -2.01 | | l lateral sulcus | |
| | 5.81 | -22.6 | -5.03 | -4.53 | | l lateral sulcus | |
| | 5.74 | -23.6 | -6.54 | -1.51 | | l lateral sulcus | |
| | 5.57 | -23.6 | -2.01 | -0.503 | | l lateral sulcus | |
| | 5.41 | -22.6 | -6.54 | 1.01 | | l lateral sulcus | |

*Table S3*. ***Significant neural clusters for OE at previous-trial-end and decision-prompt combined.***

| Cluster Index | Z | F99 coordinates x | y | z | # voxels | peak description | cluster descriptio[n] |
|---|---|---|---|---|---|---|---|
| 3 | 4.49 | 1.91E-06 | -23.6 | 18.1 | 23196 | post cingulate gyrus | cingulate and dorsomedia[l] frontal cortex |
| | 4.47 | 0.503 | -25.1 | 17.1 | | medial parietal cortex | |
| | 4.4 | 6.54 | -18.6 | 13.6 | | post cingulate sulcus | |
| | 4.37 | 1.01 | 5.53 | 11.1 | | ACC/MCC gyrus | |
| | 4.33 | 2.01 | -24.1 | 17.1 | | medial parietal cortex | |
| | 4.31 | 3.02 | -21.6 | 14.6 | | post cingulate gyrus | |
| 2 | 4.24 | 25.7 | -33.7 | -5.03 | 1917 | r inferior occ sulcus | right extrastriat[e] visual associatio[n] cortex |
| | 4.1 | 22.6 | -38.7 | 1.01 | | r inferior occ sulcus | |
| | 4.08 | 21.6 | -37.7 | 0.503 | | r inferior occ sulcus | |
| | 4.02 | 18.6 | -39.7 | -11.1 | | r lateral cerebellum | |
| | 3.97 | 17.6 | -38.2 | -9.05 | | r lateral cerebellum | |
| | 3.96 | 25.7 | -30.2 | 1.91E-06 | | r inferior occ sulcus | |
| 1 | 3.99 | 13.1 | -38.2 | 10.6 | 1833 | r visual cortex | right extrastriat[e] visual associatio[n] cortex |
| | 3.95 | 13.1 | -37.7 | 8.05 | | r second. visual cortex | |
| | 3.89 | 13.1 | -34.7 | 9.56 | | r second. visual cortex | |
| | 3.83 | 12.6 | -36.2 | 10.1 | | r second. visual cortex | |
| | 3.82 | 13.1 | -37.7 | 9.05 | | r second. visual cortex | |

| | | | | | |
|---|---|---|---|---|---|
| | 3.77 | 15.1 | -38.2 | 11.6 | r second. visual cortex |

*Table S4*. ***Significant neural clusters for ES at previous-trial-end and decision-prompt combined.***

**Supplementary Text 1**

Below we reproduce descriptions of the four different experimental designs that were used.

*Jahn et al.:*
"The task consisted of making choices between two options by responding on either the left or right touch sensor to select the left or right stimulus, respectively. A trial consisted of a given number of choices (determined by the horizon length) between these two options. Each option corresponded to one side for the entire trial. After each choice, monkeys received a reward associated with the chosen option. The reward was between 0 and 10 drops (0.5 mL of juice per drop) and was sampled from a Gaussian distribution with a standard deviation of 1.5 and mean between 3 and 7. The means of the underlying distribution were different for the two options and remained the same during a trial, such that one option was always better than the other. After each choice, monkeys also received a visual feedback on the reward. This feedback was in the form of an orange rectangle displayed in a yellow rectangular window, such that the wider the orange rectangle, the greater the amount of juice. It remained on the screen for the remainder of the trial.

At the beginning of each trial, prior to making their first choice, monkeys received 4 informative observations in total, which consisted of information about the reward they would have received if they had chosen the option. This was displayed in the same manner as reward feedback and also remained on screen during the duration of the trial. For each informative observation, a non-informative observation was presented for the other option. The non-informative observation was a white rectangle crossed by black diagonals. Half of the trials started with an equal amount of information about the two options (2 informative and 2 non-informative observations for each option), and the other half with an unequal (3 informative and 1 non-informative observations). The order and side were randomly determined.

A critical parameter was the number of choices in each trial (horizon length). In **short horizon** trials, monkeys were only allowed 1 choice before a new trial with new stimuli started, whereas in **long horizon** trials, they were allowed to make 4 choices between the options. Horizon conditions were blocked (5 consecutive trials of equal horizons) and alternated in the session. A second key manipulation was whether feedback was received only for the option they chose (**partial feedback** condition) or whether they received information about both the reward they received for the chosen option *and* the reward they would have received for selecting the alternative option (**complete feedback** condition).

A trial would proceed as follows: After an inter-trial interval during which the screen was black, the stimuli were displayed, consisting of a large grey rectangle and the 4 horizontal bars of feedback information. The length of the grey rectangle corresponded to the length of the horizon, which each line corresponding to a choice, simulated or actual. Informative or non-informative stimuli were displayed on the first four lines. After the display of the stimuli, a red dot at the center of the screen disappeared, and monkeys were then allowed to choose between the two options by touching the corresponding sensor (in less than 5,000 ms or the trial restarted). A red rectangular frame appeared around the line on the side of the chosen option. After a delay, the outcome—the reward feedback—was displayed inside the rectangle. In complete feedback condition only, the reward that would have been gained on the other side (informative stimulus) was also displayed at the same moment. After an additional delay, a white star appeared on the screen, and the reward was delivered. After the end of the reward delivery, the star disappeared. In short horizon blocks, a new trial started after the inter-trial interval delay. In long horizon trials, the red dot appeared and then monkeys could choose among the options. The events leading to the reward were similar than for the first choice, but the delays were shorter. At the end of the fourth choice, a new trial started. The feedback condition monkeys were in was not explicitly cued but instead fixed both within and across several sessions (6 to 10 consecutive sessions). Sessions after a switch from one feedback condition to the other were included in the analysis since it only took one choice for monkeys to know the feedback condition."

*Grohn et al.:*

"Each trial began with a blank screen (2–4 s, mean 3 s) followed by a presentation of the rectangle. The monkeys responded by touching either of 2 custom-built infrared sensors placed in front of them. Each manual response was classified as either correct (the monkey touched the response sensor adjacent to the stimulus) or incorrect (the monkey touched the other sensor). Each correct response yielded a juice reward of 1, 2, or 3 drops (approximately 1.5 ml each drop) after a delay of 200 ms. The juice delivery took 1.5 s, and after an intertrial interval of 2–4 s (mean 3 s), the next trial began (Fig 1A). If the response was incorrect, the trial was repeated until the monkey made the correct response. Importantly, the spatial cue position (left or right) varied independently of reward magnitude (1, 2, or 3 drops) that was given for each correct trial. Reward was thus decorrelated from spatial position of the stimulus.

The task design enabled us to examine VS because the side on which the stimulus was shown reversed after 11–19 trials on the same side (mean 15 trials) although occasional stimuli appeared on the opposite side throughout. Eleven to 13 sessions of 150 rewarded (i.e., correct) trials were collected for each of 6 animals while they were in the MRI scanner. Thus, we could examine VS by comparing trials on which the stimulus had appeared on the same or the opposite side compared to the previous trial. Three types of sessions were performed by the monkeys, as follows.

**Stable/Unlearnable sessions**

In 6 sessions, 1 and 3 drops were delivered randomly in 90% of the reward trials (45% for each reward size), and 2 drops were delivered in 10% of rewarded trials. Mean reward over a session was kept at 2 drops. However, even if 2-drop rewards accorded with the average reward expectation, they were only rarely delivered, and so they were in this sense the most surprising outcomes. Therefore, it was possible to identify activity related to RRE by comparing 2-drop reward outcomes with all other outcomes, and it was possible to identify sRPE-related activity by calculating the parametrically varying sRPE associated with each outcome. The sRPE and RRE regressors shared only 0.049% of variance. One weakness of the schedule, however, is that it is static and does not change over the course of the session. Because no learning is possible in such situations, learning mechanisms may not be deployed. We attempted to remedy this deficiency by using additional schedules.

**Changing/learnable sessions**

In 2 different reward schedules (each comprising 2 sessions), the mean reward changed either from an average of 1.5 drops to an average of 2.5 drops or in the opposite direction, from 2.5 drops down to 1.5 drops halfway through the session. In these changing/learnable sessions, either 1 or 3 drops were delivered on 90% of trials on average, across the whole session. Therefore, once again it was possible to identify activity related to RRE by comparing 2-drop reward outcomes with all other outcomes, and it was possible to identify sRPE-related activity by calculating the parametrically varying sRPE associated with each outcome. However, it is possible that effects may be stronger in the changing/learnable sessions than the stable/unlearnable sessions because the reward environment is genuinely getting either better or worse. In order to estimate which is the case, animals must pay attention to outcomes. In these sessions, the sRPE and RRE regressors shared 0.1551% of the variance.

**Equiprobable sessions**

In a third, control condition (comprising 2 sessions), we kept the average reward stable, but each reward magnitude had an equal probability of 1/3, thereby eradicating any reward frequency effects.

Therefore, once again it was possible to identify sRPE-related activity by calculating the parametrically varying sRPE associated with each outcome. Now, however, there may be less RRE effect because 2-drop reward outcomes are no less frequent, and therefore no more surprising, than 1- or 3-drop outcomes."

*Bongioanni et al.:*

"Experiment 1 included 12 fMRI sessions of 180 binary choice trials per subject; no-response trials were repeated at the end of each session. Trials were divided into six conditions with 30 trials each. Factors were:

familiarity of the options:

◦ familiar (both options were familiar)

◦ novel (both options were novel)

dimensionality of the pair:

◦ consistent (one option was associated with both higher reward magnitude and higher reward probability than the other option)

◦ one-dimensional (the two options had one identical attribute, but differed in the other attribute)

◦ inconsistent (one option had higher magnitude and the other option had higher probability)

Condition order across trials was pseudo-randomized. Investigators were not blinded to allocation during experiment and outcome assessment. The expected value range of the stimuli (that is, magnitude times probability) during training was 0.5 to 6 drops; this was preserved during testing. Combinations with an expected value outside the familiar range were excluded.

Average expected value sum (left value plus right value) was matched across all conditions (6 ± 0.02 drops). Value difference was necessarily different across dimensionality conditions; but was matched across familiarity conditions, both in terms of mean and variance (within ranges of 0.02 drops and 0.035 squared drops, respectively). Finally, the correlation between the best value and the worst value offered across trials was lower than $r = 0.33$ when merging all familiar conditions and when merging all novel conditions. This stimulus schedule was designed to allow the dissociation of neural signals related to the comparison signal from the value sum.

For each trial, stimuli were presented for up to 60 s until response (median response time (RT) = 952 ms). The action-outcome delay lasted 3.5–4.5 s (uniform distribution). A visual cue indicated either positive (upward triangle) or negative (downward triangle) outcome; if positive, juice was simultaneously delivered through a spout. The outcome cue lasted 3 s whether positive or negative. The inter-trial interval (ITI) had a duration of between 5 and 7 s (uniform distribution) (Fig. 2a).

These delays made it possible to decorrelate the BOLD signal related to decision-making and response from the BOLD signal related to subsequent feedback processing and juice consumption. The haemodynamic response function (HRF) in the macaque is faster than in humans34, peaking in approximately 3 s. Motor response-related activity was accounted for by confound regressors in the GLM analysis as described below."

*Khalighinejad et al.:*

"At the beginning of each trial an empty frame (8 × 26 cm) appeared on the left or right side of the screen. The frame gradually filled with dots (round circles, r = 0.3 cm, max number of dots = 25) emerging from top to bottom. Animals could terminate the trial, at a time of their own choice, by touching a custom-made infra-red touch sensor, on the side corresponding to the image. The trial continued if they touched the opposite side. The probability of getting reward increased as more dots appeared on the screen, following a sigmoid curve. The probability distribution was drawn from a sigmoid function. The input to the function was a vector corresponding to the number of dots from 1 to 25. The midpoint of the curve was at dot #12 (50% chance of getting reward) with the steepness of 0.5. The probability distribution was constant across the trials and the sessions. The color of the frame and dots varied from trial to trial but remained constant within a trial. The color indicated potential reward magnitude and could be red, green or blue, indicating one, two or three drops of juice, respectively. In addition to the color, the speed of the dots appearance also varied from trial to trial. A new dot appeared every 100, 200 or 300 ms. Animals had the option to respond, any time from the beginning of the trial (appearance of the empty frame) to 300 ms after the frame was filled (appearance of the last dot). If they responded, they were offered drops of juice or no juice, based on the probability distribution at the time of response. There was a delay of 4 s between response and outcome (action-outcome delay). Successful and unsuccessful outcomes were indicated by an upward and downward pointing triangle, respectively. The triangle remained on the screen for 2 s. If rewarded, drops of blackcurrant juice were delivered by a spout placed near the animal's mouth during scanning. Each drop was composed of 1 mL blackcurrant juice. No juice was delivered when the trial was not rewarded. After the outcome phase, they proceeded to the next trial after a 3, 5 or 7 s inter-trial interval (ITI). ITI varied in blocks of 30 trials in a pseudo-randomized order. Specific patterns on the left and right side of the screen indicated the ITI block. If animals did not respond by 300ms after the emergence of the last dot, the frame disappeared, and they had to wait for 4 s (equivalent to action-outcome delay) + 3, 5 or 7 s (ITI) for the next trial to start. Animals were given 40min to perform the task at each session. The task finished after 40 min, regardless of the number of trials done. Each animal performed ten to twelve sessions in the MRI scanner. The experiment was controlled by Presentation software (Neurobehavioral Systems Inc., Albany, CA)."
